# Leveraging Distributed Biomedical Knowledge Sources to Discover Novel Uses for Known Drugs

**DOI:** 10.1101/765305

**Authors:** Finn Womack, Jason McClelland, David Koslicki

## Abstract

Computational drug repurposing, also called drug repositioning, is a low cost, promising tool for finding new uses for existing drugs. With the continued growth of repositories of biomedical data and knowledge, increasingly varied kinds of information are available to train machine learning approaches to drug repurposing. However, existing efforts to integrate a diversity of data sources have been limited to only a small selection of data types, typically gene expression data, drug structural information, and protein interaction networks. In this study, we leverage a graph-based approach to integrate biological knowledge from 20 publicly accessible repositories to represent information involving 11 distinct bioentity types. We then employ a graph node embedding scheme and use utilize a random forest model to make novel predictions about which drugs can be used to treat certain diseases. Utilizing this approach, we find a performance improvement over existing computational drug repurposing approaches and find promising drug repositioning targets, including drug and disease pairs currently in clinical trials.

## 1. Introduction

The collection and curation of large repositories of information has led to significant advancements in the training of machine learning approaches in areas as diverse as image recognition [42,89], textual analysis [47, 63, 98], and speech recognition [45, 77, 83, 102]. In particular, in the realm of biomedical sciences, large relatively mature collections of information have been assembled covering areas such as genes and proteins [12,58], biological processes and pathways [5,57,71], drugs [1,46,76], and diseases [53,80,85]. Repositories such as these, with varying degrees of success, have previously been used to investigate the problem of drug repurposing [75,97], also called drug repositioning, the main subject of this manuscript. Drug repurposing is the process of using a previously medically approved drug to treat a disease different from than the ones originally intended. For example, the drug thalidomide was originally developed in the 1950’s to be an anti-anxiety medication [70], but fell out of favor due to its causing abnormal physical development. However, it was subsequently discovered that thalidomide can be used to treat certain forms of cancer [3]. Examples such this are bolstered by the economic advantage of drug repurposing: utilizing a well understood, well-characterized drug that has already been approved for some treatments reduces the likelihood that clinical trials will fail due to safety concerns, and the average $1.3B cost of bringing a new drug to market is reduced to $8.3M when repurposing an existing drug [78].

While machine learning approaches to the drug repurposing problem have been proposed [16, 17, 20, 21, 22, 56, 67], these methods typically use only a few sources of information, and typically only a few characteristics of a given drug and disease, such as genomic information, chemical structure information, and/or drug interaction information [65]. More recently, integrative approaches that combine information from multiple data sources have been proposed which were shown to out-perform non-integrative approaches [75, 101]. However, even in these integrative approaches, only three [75] or four [101] different data sources are integrated into a machine learning approach.

In this study, we investigate the utility of integrating a variety data types (11 in total) from multiple data sources (20 in total) in order to enhance the accuracy of computational drug repurposing. Figure 1 describes the high-level approach we take to this problem. Briefly, different data are represented as nodes in a graph (here called a “knowledge graph”) and data sources (here called “knowledge sources”) are used to connect nodes in the graph with edges/relationships and annotate nodes and edges with additional information. The resulting knowledge graph we created possesses over 124K nodes and 7.6M edges. We then apply a graph node embedding algorithm that learns low-dimensional, neighborhood preserving, vector representations for nodes in the knowledge graph. After curating training data, we then design a random forest to learn which drugs treat which diseases and generate novel drug repurposing candidates.

**Figure 1.**
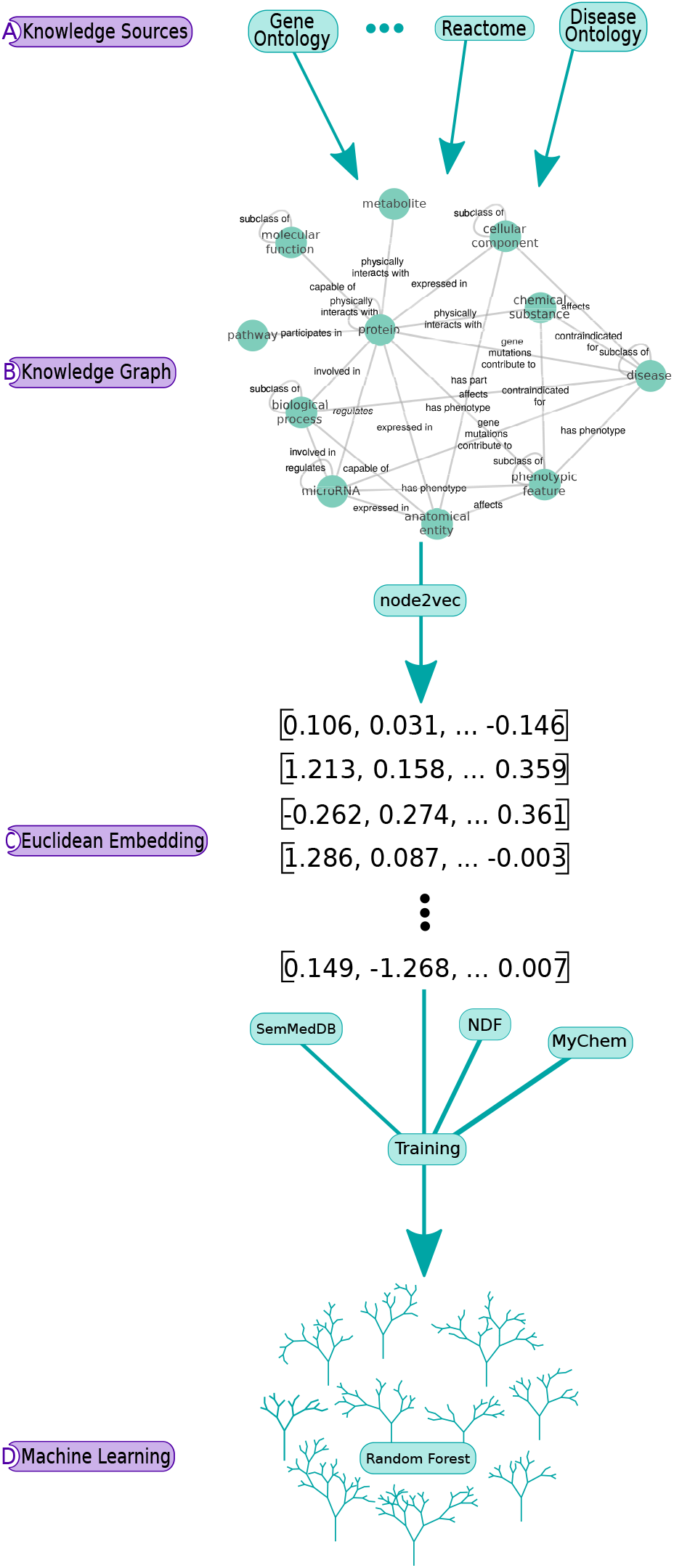
Overview of the work flow. Part A) depicts how we query external repositories of information to populate B) a graph database. Part B) depicts the node bioentity types and the relationships that connect them (given in more detail in Tables 1 and 2). The actual knowledge graph possesses over 124K nodes and 7.6M relationships. In part C), node2vec is used to generate Euclidean embeddings of drug and disease nodes which are then combined with true positive/negative training examples populated from additional public repositories and are used to train D) a random forest classifier.

## 2. Methods

The approach we take is depicted in Figure 1: we begin in part A) of Figure 1, where a variety of biomedical repositories are queried to populate B) a graphical database, referred to as a “knowledge graph”, that describes the relationships between the various bioentities. The graph node embedding method node2vec [64] is then used to generate vector representations of each node in the graph corresponding to a “drug” or “disease” node type, as depicted in part C). We then utilize three additional repositories to generate training data: drug-disease pairs for which it is known that the drug either treats/is indicated for the disease (a true positive case) or does not treat/is contraindicated for the disease (a true negative case). In part D), this is used to train a random forest model in order to predict drug-disease pairs for which the drug is predicted to treat the given disease. In the following, we outline the details of our approach.

**Table 1.** List of nodes in the knowledge graph. Each node is assigned a label, given in the first column, indicating what type of bioentity it represents. The second column gives the count of such nodes in the knowledge graph, and the last column specifies which knowledge sources were queried to populate these nodes in the knowledge graph.

| Node Bioentity Label | Node Count | Knowledge Sources |
| --- | --- | --- |
| Biological process | 30716 | GO [6, 39] |
| Protein | 20514 | UniProtKB [40] |
| Disease | 19573 | DOID [59], OMIM [53], MONDO [71] |
| metabolite | 18145 | KEGG [58] |
| Molecular function | 12140 | GO [6, 39] |
| Phenotypic feature | 12089 | HP [62] |
| Cellular component | 4375 | GO [6, 39] |
| Chemical substance | 2226 | CHEMBL [10, 46] |
| Pathway | 2215 | Reactome [41] |
| microRNA | 1697 | NCBIGene [68] |
| Anatomical entity | 1121 | UBERON [72], CL [84] |
| **Total:** | 124,811 |  |

**Table 2.** List of relationships/edges in the knowledge graph. Each relationship is assigned a type, given in the first column, indicating the kind of relationship it represents. The second column gives the count of such edges in the knowledge graph, and the last column specifies which knowledge sources were queried to populate these edges in the knowledge graph.

| Relationship Type | Edge Count | Knowledge Source |
| --- | --- | --- |
| Participates in | 1,558,894 | Reactome [41] |
| Has Phenotype | 1,512,732 | BioLink [11] |
| Physically interacts with | 1,478,924 | KEGG [58], UniProtKB [40], ChEMBL<br>Pharos [76], Reactome [41], PC2 [18] |
| Regulates | 1,477,708 | miRGate [4], PC2 [18], GeneProf [50, 51] |
| Expressed in | 828,976 | BioLink [11], GO [6, 39] |
| Involved in | 275,946 | GO [6, 39] |
| Subclass of | 201,314 | GO [6, 39], Monarch SciGraph [71],<br>DOID [59] |
| Capable of | 125,526 | Monarch SciGraph [71], GO [6, 39] |
| Gene associated with condition | 79,238 | BioLink [11], DisGeNet [79] |
| Contraindicated for | 42,970 | MyChem.info [94] |
| Gene mutations contribute to | 11,538 | OMIM [53] |
| Indicated for | 8,940 | MyChem.info [94] |
| Affects | 6,130 | Monarch SciGraph [71], BioLink [11] |
| Has part | 34 | Monarch SciGraph [71] |
| **Total:** | 7,608,870 |  |

### 2.1 Knowledge graph construction

We queried 20 separate, publicly available repositories of information via RESTful API calls in order to populate a Neo4j [92] graphical database with 11 different types of bioentities. The node bioentity types, counts, and originating knowledge sources are given in Table 1. In total, 124,811 different nodes were created in the knowledge graph. The knowledge sources also provide relationships between bioentities, and these were used to form edges between bioentity nodes in the knowledge graph. Table 2 lists the various relationships that were obtained from the knowledge sources. From a very abstracted perspective, each of the knowledge sources can be thought of as containing information (curated either manually or computationally) about the respective bioentities contained in that repository. In total, 7,608,870 edges were populated in the graph. The code used to generate this knowledge graph is available at: https://github.com/RTXteam/RTX/tree/reasoning_lists/code/reasoningtool/kg-construction and a static Neo4j dump of the resulting Knowledge Graph is available at: http://kg1endpoint.rtx.ai/20180807-000318.tar.gz.

### 2.2 Node embedding

After constructing the knowledge graph in Section 2.1, we then applied a machine learning package called node2vec [48] from the SNAP Library [64] which learns the structure of the graph using random walks and generates a feature vector of fixed length for each node in the graph. This method has several parameters that can be tuned in order to change the resulting vector embedding. The parameters we focused on where the two hyperparameters *p, q*, as well as the dimension of the embedding *d*, the length of the random walk *l*, the number of random walks taken *r*, and the number of epochs in the stochastic gradient descent subroutine *e*. To determine the best set of parameters we ran a grid search and compared the resulting mean F1 scores from 10-fold cross-validation. Using the model specified in 2.4, we obtain optimal parameters of *p* = 1, *q* = 5, *d* = 256, *l* = 300, *r* = 15, and *e* = 5.

### 2.3 Training data generation

We generated our training data using 3 different knowledge sources to obtain true positive/negative drug and disease/phenotypic feature pairs as detailed in Table 3. The true positive training data consists of drug and disease (or phenotypic feature) pairs of nodes for which it is known that the drug treats or is indicated for the disease/phenotypic feature.

**Table 3.** List of sources used to generate training data. The second and third columns give the counts of true positives (indications) and true negatives (contraindications or no effect) respectively.

| Source | Positives (Treats) | Negatives (Not Treats) |
| --- | --- | --- |
| SemMedDB [60] | 73,976 | 2,012 |
| MyChem [94] | 3,673 | 23,606 |
| NDF-RT [15] | 3,111 | 5,586 |
| **Total:** | 80,760 | 31,204 |

True negative data consists of drug and disease/phenotypic feature nodes for which it is known the drug either does not treat or else is contraindicated for the disease.

The first knowledge source, SemMedDB [60] is a repository of subject-predicate-object triplets extracted from all PubMed [2] abstracts as of December 31, 2017 via SemRep [82], a semantic interpreter of biomedical texts. The SemMedDB predicates “Treats” and “NEG Treats” were used to partially generate positive and negative training data respectively, which were then mapped to drug and disease (or phenotypic feature) tuples in the knowledge graph. We selected only those tuples which occurred three or more times in SemMedDB to ensure sufficient literature support.

We also used the MyChem API [1, 94] to find indicated diseases/phenotypic features (true positive) and contraindicated diseases/phenotypic features (true negative) for the drugs in our knowledge graph. MyChem is a BioThings [95] high-performance web service that aggregates a variety of public repositories of information on chemical substances.

Lastly, we also utilized The National Drug File - Reference Terminology [15, 90] which is an ontology produced by the U.S. Department of Veterans Affairs, Veterans Health Administration to check drug interaction, indications, and contraindications. As before, we extracted (via Protege [74]) the indicated diseases/phenotypic features and contraindicated diseases/phenotypic features for the drugs in our knowledge graph.

Finally, we took the vector embeddings (see Section 2.2) of each training example drug and disease and combined them with the Hadamard product to generate a feature for each drug and disease pair. In total, 80,760 true positive and 31,204 true negative training examples were generated, representing 3,192 distinct diseases/phenotypic features and 1,571 distinct drugs.

### 2.4 Model specification and performance

We evaluated the performance of both a logistic regression model, a kernel support vector machine model, and a random forest model to discriminate our training data, judging performance via mean F1 score and area under the receiver operating characteristic (ROC) curve (AUC) both generated with 10-fold cross-validation. To prevent information leakage due to drug similarity, we ensured that in each cross-validation fold, no drug classes were shared between the validation set and the training set. Drug classes were determined using the MyChem API [1].

Importantly, while the training data reflects a bias towards publishing positive results (the training data has roughly 2.6 times more positive examples than negative examples), it is much more likely that a randomly selected drug will not treat a randomly selected disease. Thus, in addition to the F1 score and AUC, we also calculate the percentage of randomly selected drugs and diseases that were predicted to be positive examples (i.e. “drug treats disease”) as a function of the classification threshold. Such a figure is given in Figure 3. Thus, an optimal model would not only have a high F1 score and AUC, but would predict that it is highly unlikely that a randomly selected drug will treat a randomly selected disease.

**Figure 2.**
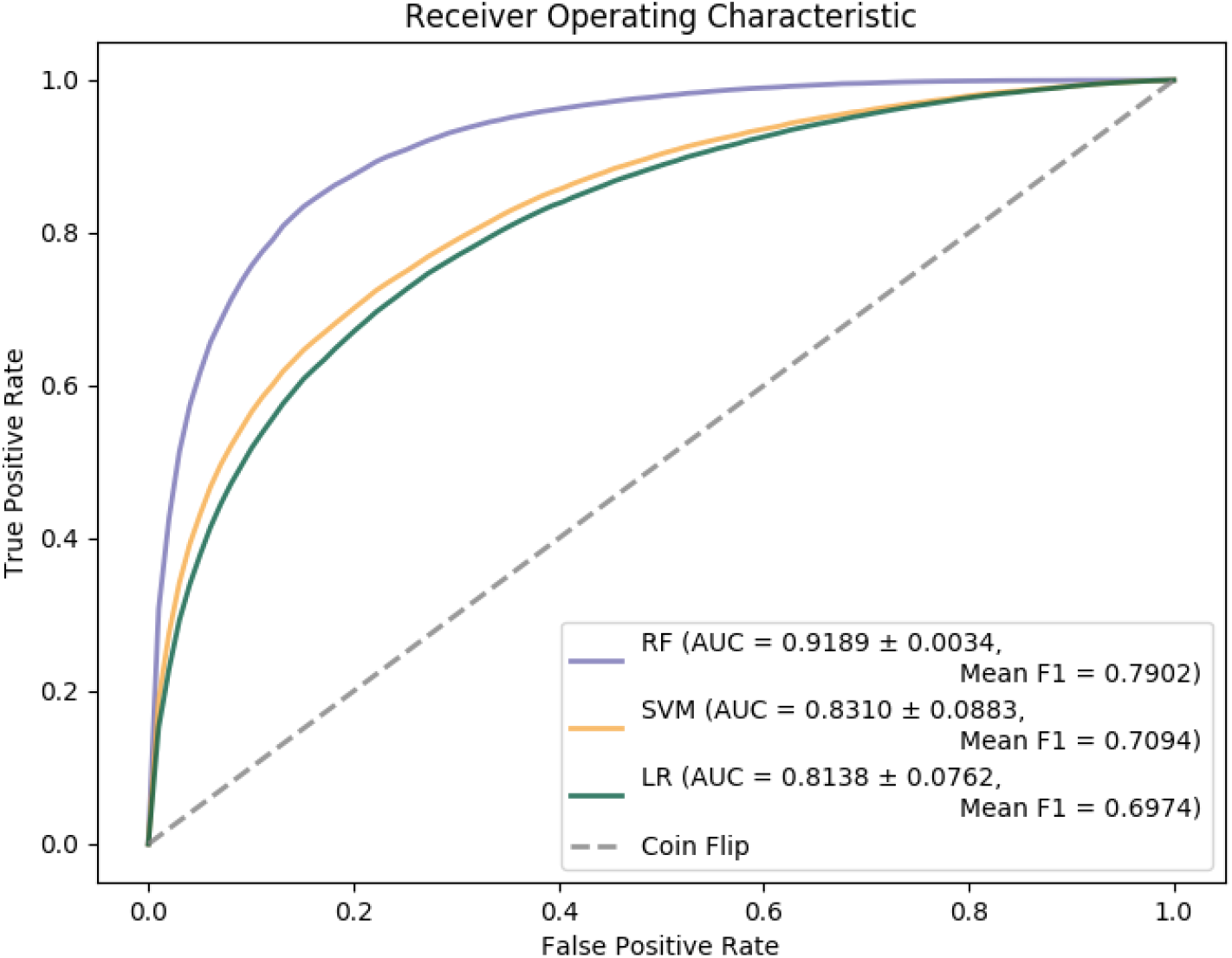
ROC curve comparing random forest, support vector machines, and logistic regression models demonstrating the superior performance of the random forest approach. The plot was generated with 10 fold cross-validation while grouping drugs of the same class into the same fold. The inset displays the mean AUC and and F1 scores. The random forest model uses a maximum depth of 29 with 2,000 trees.

**Figure 3.**
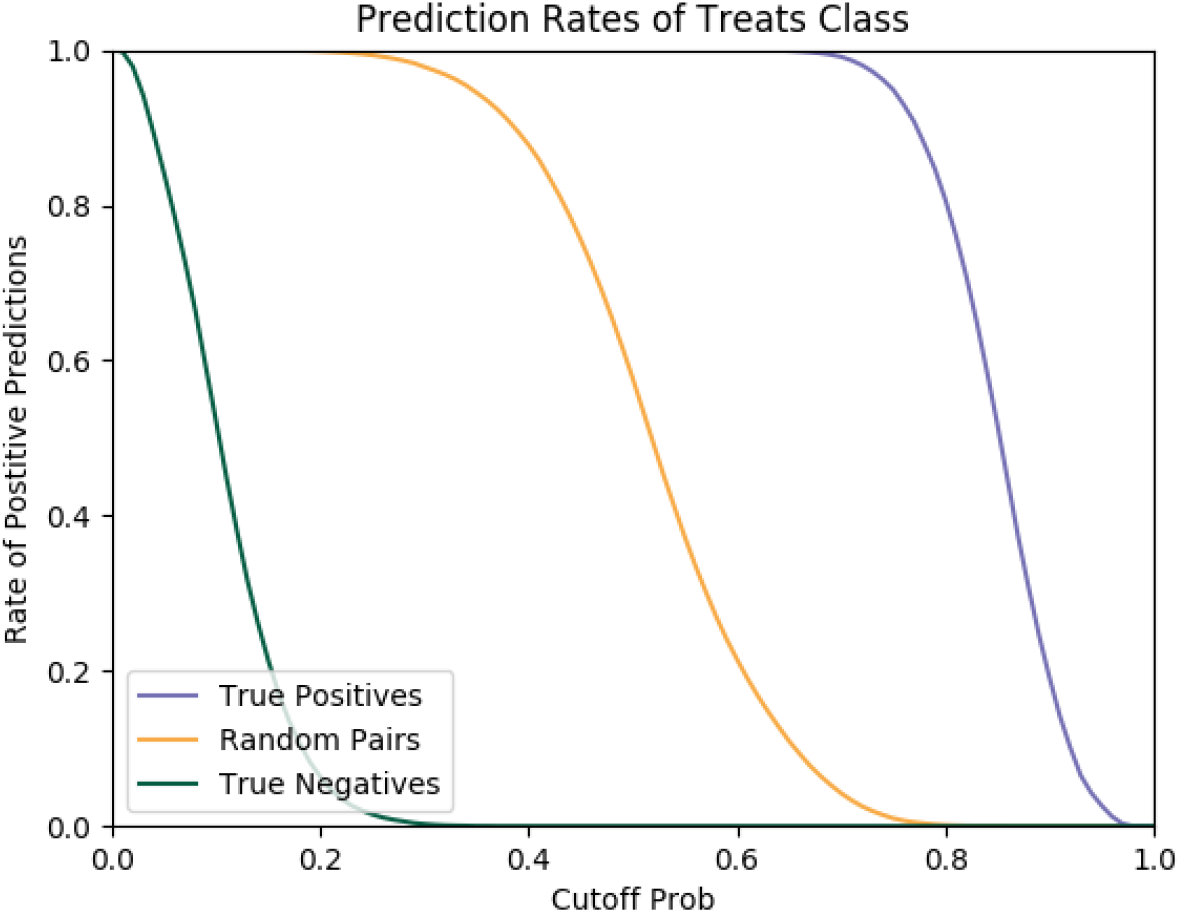
Plot depicting the percentage of randomly selected drugs and diseases that were predicted to be positive examples as a function of the classification threshold while using the optimal random forest model. Blue and green lines indicated prediction rates for true positive and true negative classes respectively, while the yellow line is calculated using 100,000 random pairs of drugs and diseases.

To combat the negative effects of the training data class imbalance [19], in the random forest training, we weighted each class inversely proportional to its frequency. We also determined that when using 2,000 trees, the depth of 29 was the first depth that was within 4% of the minimum out of bag error over all depths up to depth 40.

Hence by using these criterion, we found that a random forest utilizing 2,000 trees and a maximum depth of 29 maximized the F1 score, AUC, retained a majority of the true positives while assigning low probability of “treats” to a majority of the randomly selected drug and disease pairs, and is also unlikely to be over-fitting the training data. In comparison, both a logistic regression and support vector machine approach suffered in terms of F1 score and AUC (see Figure 2).

This random forest model had a mean F1 score of 0.7902 and mean AUC of 0.9189 using 10-fold cross validation, as shown in Figure 2. As depicted in Figure 3 using the random forest model, for a classification threshold of 0.75, 98.995% of the randomly selected drugs are not predicted to treat randomly selected diseases, while still correctly classifying 94.889% of the true positive training examples. Hence, according to 3, we can be confident that a positive “treats” prediction for a drug disease pair above 75% is likely not to be a false positive.

## 3. Results

### 3.1. Validation of results

Table 4 contains an assortment of drugs predicted to treat certain diseases by the model specified in Section 2.4. This list was generated by looking at a particular disease/phenotype, generating treatment probabilities according to the model, and then selecting the top few drugs according to these probabilities. Importantly, these drug/disease pairs were not included in the training data and represent novel predictions. For a few of the examples contained in Table 4, we present here additional external evidence to support the validity of the results obtained by our machine learning model.

**Table 4.** A selection of high-probability drug and disease pairs as predicted by our model. The first and second columns indicate the drug and disease/phenotype pairs, while the third column provides literature support for the treatment of the disease by the drug. The fourth column indicates if `clinicaltrials.gov` provides information about ongoing clinical trial for the treatment of the disease by that drug. The last column indicates the treatment probability as returned by our random forest model. None of these examples were included in the training data.

| Drug | Disease/Phenotype | Documentation | Clinical Trial | Treat Prob |
| --- | --- | --- | --- | --- |
| Valproic acid | Alzheimer’s disease | [28, 38, 96, 99] | Phase 3 | 0.873597 |
| Anakinra | Ankylosing Spondylitis | [49, 87] |  | 0.982593 |
| Secukinumab | Ankylosing Spondylitis | [7, 13, 29] | Phase 4 | 0.949114 |
| Teriparatide | Arthritis | [23, 43] | Phase 4 | 0.885561 |
| Ixekizumab | Arthritis | [24, 69] | Phase 4 | 0.849326 |
| Azithromycin | Bronchitis | [8, 26, 31, 36] | Phase 4 | 0.816302 |
| Prednisone | Bronchitis | [81] |  | 0.789305 |
| Interferon alfa-2b | HIV | [33] | Phase 2 | 0.940958 |
| Peg-interferon alfa-2b | HIV | [25, 32] | Phase 3 | 0.907307 |
| Oxprenolol | Hypertension | [100] |  | 0.842847 |
| Rituximab | Osteoporosis | [54] |  | 0.902776 |
| Anakinra | Posterior Uveitis | [9, 88] |  | 0.957913 |
| Cyclosporine | Posterior Uveitis | [9, 34, 73] | Phase 3 | 0.938793 |
| Dexamethasone | Posterior Uveitis | [30, 66] | Phase 4 | 0.838129 |
| Deferoxamine | Tendonitis | [44] |  | 0.957913 |
| Tocilizumab | Type I diabetes mellitus | [27] | Phase 2 | 0.825334 |
| Doxycycline | Type II diabetes mellitus | [35, 37, 91] | Phase 4 | 0.905237 |

Among these, the drug dexamethasone is predicted to treat posterior uveitis (a type of inflammation in the eye) with a probability of 0.838129. Using an additional knowledge source, Colombia Open Health Data (COHD) [86] which provides anonymized key word search over 1.7 million health records, we find that dexamethasone co-occurs with posterior uveitis at a rate 2.789 times more frequently than would be expected in a general population. This may indicate that dexamethasone is being prescribed as an off-label treatment for this disease. The validity of this result is bolstered by a phase 3 clinical trial which found that dexamethasone is an effective treatment for treating posterior uveitis [66], and the recent recommendation by the United Kingdom’s National Institute for Health and Care Excellence to use dexamethasone for the treatment of uveitis.

The use of secukinumab for the treatment of ankylosing spondylitis (an inflammatory arthritis of the spine and joints) was found to have a treat probability of 0.949114 and using COHD we find that secukinumab co-occurs with ankylosing spondylitis at a rate 5.942 times more frequently than would be expected in a general population. Secukinumab was shown to reduce the signs and symptoms of ankylosing spondylitis in both long [13] and short term [7] phase 3 studies. A phase 4 clinical trial is also currently recruiting to study different doses of secukinumab as a treatment for ankylosing spondylitis [29].

The use of doxycycline for the treatment of type II diabetes mellitus was found to have a treat probability of 0.905237. Doxycycline has been shown to decreases systemic inflammation and improve glycemic control in low doses in diabetic mice [91]. Additionally, a phase 4 clinical trial was performed to assess if doxycycline will enhance insulin sensitivity and decreases inflammation in obese participants with type 2 diabetes [35].

The use of valparic acid for the treatment of Alzheimer’s disease was found to have a treat probability of 0.873597. Valparic acid has been hypothesised to help with Alzheimer’s disease as it can induce neurogenesis of neural stem cells [99] and it has been shown to be a potential therapeutic approach for Alzheimer’s using mouse models [96]. Additionally, a phase 3 clinical trial has been preformed for the treatment of Alzheimer’s with valparic acid [38] as well as a phase 1 clinical trial exploring it as a potential preventive medicine [28].

Lastly, the use of ixekizumab for the treatment of arthritis was found to have a treat probability of 0.849326. A phase 3 clinical trial found that the American College of Rheumatology response criteria 20 was achieved after 12 and 24 weeks [69], thus demonstrating effectiveness over a placebo. There is also an active phase 4 clinical trial studying the efficacy of ixekizumab on arthritis [24].

## 4. Comparison to existing approaches

### 4.1 Overview of existing approaches

Here, we will compare and contrast the current approach to several other approaches to *in silico* drug repurposing. Previously, methods generally focused on the guilt-by-association principle, that structurally similar drugs will have similar targets, which can be leveraged to determine novel drug-target pairs. In particular, identifying physical structural similarities between drugs or drug receptors which allow for prediction of binding affinities [56]. These include docking based screening techniques which use computational molecular models of physical docking mechanisms to predict the fitness of drug-target pairs. Such methods require detailed information regarding the mechanisms of interaction and the systems involved [22]. Alternate similarity metrics, such as based in side effect overlap [16], have also been utilized to determine unexpected drug-drug similarities and interactions.

Many recent approaches utilize machine learning techniques to predict drug-target interactions. Following in the vein of the structural similarity based methods described above, feature sets generated from concatenating hashed chemical descriptors for drugs and protein targets analyzed by random forest regression have achieved AUC’s of 0.96 [17]. Machine learning techniques have also been put forth in the context of network-based inference of drug-target interaction. These methods utilize graph-based data structures to predict novel candidates for drug repurposing, under a variety of schemes for feature generation. Utilizing a bipartite graph consisting of drugs and targets, edges indicating known interactions populated from the DrugBank database [93] and a simple scheme of two-step diffusion to infer similarity, Cheng et al. showed that a network based inference system outperformed various structural similarity based inference methods [21].

In these previous approaches, a restricted number of data sources are typically utilized (such as drug structure and side effects, or chemical descriptions and protein targets). More recent approaches, including the one presented in this manuscript, have sought to integrate a variety of heterogeneous data sources in a network based model. Chen et al. [20] utilized a heterogeneous network composed of weighted drug-drug similarities, weighted protein-protein similarities and and known drug-target interactions. Using the weights from these data sources, they constructed a set of transition probabilities for a random walk on each of the drug and target networks, as well as transition probabilities between networks. By fixing parameters for the transition between networks for a random walk and the probability for restart in the random walk, they then numerically determined stationary probabilities for the random walk beginning from a fixed drug. From these stationary probabilities they defined the the most likely novel drug-target interactions between the fixed drug in question by ranking via the probability of find a random walker at a given target. This differs from the method described here in several ways: the much smaller number of data sources used, the lack of integration between networks and the necessity of evaluating drugs individually for potentially novel interactions.

More recently, Luo et al. [67] constructed a computational pipeline for predicting drug-target interactions called DTINet which similarly uses a heterogeneous set of networks of biologically significant entities containing some 12K nodes and 1.9M edges, built from known interactions between drugs, proteins, diseases and side-effects. Similar to the method presented in this manuscript, low-dimensional feature vectors were generated from this network via random walk with restart, which approximates the network diffusion. This was performed for each of the networks examined independently, followed by diffusion component analysis, which acts a dimension reduction and denoising scheme. This builds a single set of features for drugs and targets from the set of networks. Determination of predicted drug-target interactions is done by solving for a linear projection of drug features onto target features which minimizes the regularized *L*_2_ discrepancy from the binary matrix of known drug-target interactions. The predicted novel interactions are then between those drugs which are geometrically close to targets under this projection. Use of this method led to the prediction of a set of novel COX inhibitors, results which were supported by the researchers use of *in silico* docking modeling methods. This method differs most substantively from the method presented here in that again the networks are constructed separately, before being merged into a single set of feature vectors via diffusion component analysis.

As an approach most similar to the one presented in this manuscript, Himmelstein et al. [55] constructed a knowledge graph consisting of 47K nodes of 11 data types interconnected by 2.2M edges of 24 types for use in drug repurposing. This knowledge graph was constructed from 29 public resources, giving information regarding relationships between drugs, diseases, genes, anatomies, pathways, biological processes, molecular functions, cellular components, pharmacologic classes, side effects, and symptoms. Initial features for training a logistic regression model were generated via a prior probability for efficacy determined from drug and disease connectivity, node degrees for a subset of edge types and a set of degree weighted path counts (DWPC), which inversely weights path counts of a fixed node and edge composition between a drug-disease pair by the degree of intervening nodes. This set of features was reduced by the use of a cross-validated elastic net to a set of 31 features. Their model was trained using 755 known true positives and 29,044 known true negatives. Using this method they were able to, for example, identify clinically significant treatments for nicotine addiction and epilepsy.

Given the similarity of the currently proposed approach to that of Himmelstein et al., we chose this method to carry out a comparison based on performance characteristics. We will refer to the approach of Himmelstein et al. as DWPC given the importance of the degree weighted path counts in their approach. We will refer to the method presented in this manuscript as the node2vec approach given that we use node2vec for feature generation.

### 4.2 Comparison to DWPC method

In order to perform a fair comparison, we performed the DWPC approach as detailed in [55], but applied it to the larger knowledge graph described in Section 2.1. We utilized the same set of 147 diseases evaluated in [55] and all 2,226 drugs for both approaches. We utilizing the same 10-fold cross validation described in Section 2.4 and the optimized random forest model described in Section 2.4. As seen in the ROC curve given in Figure 4, performance was nearly identical between the node2vec and DWPC approaches on this reduced training set, with the node2vec approach only slightly improving upon the AUC and F1 metrics, with a mean improvement of 1.4% and 4.0% respectively over DWPC.

**Figure 4.**
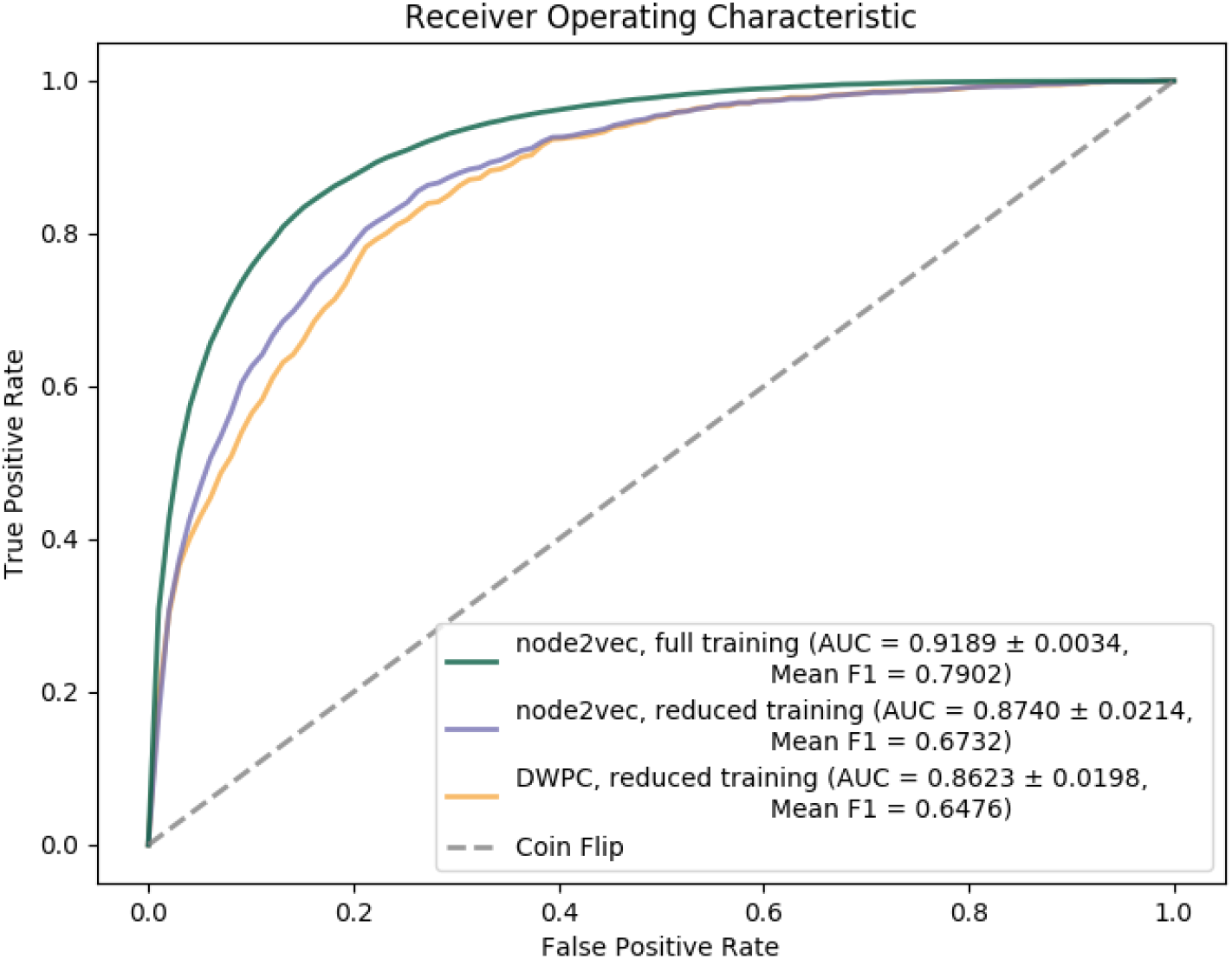
ROC curve using the knowledge graph generated according to Section 2.1, comparing the Himmelstein et al. degree weighted path count method (DWPC) with the node2vec approach we took. We used the same 10-fold cross validation and optimized random forest model as before. The green line indicates the node2vec approach trained on all 3,192 distinct diseases/phenotypic features. The blue and yellow lines indicate the performance of the node2vec and DWPC approaches respectively when restricting the training set to only 147 diseases to facilitate a direct comparison due to efficiency constraints experienced by the DWPC method.

However, with respect to computational efficiency, we find significantly different performance characteristics. To quantify this, on the same 24 core/48 thread, 2.4 GHz, 256 GB server, we ran the DWPC approach with a varying number of diseases along with the node2vec approach using the full training data described in Section 2.3. The DWPC approach requires direct access to the knowledge graph, and this was hosted locally in neo4j. Figure 5 depicts the run-time performance of both algorithms as measured by wall time. Included in Figure 5 is a linear line of best fit and extrapolation depicting that it would require 5,549.29 hours for the DWPC approach to run on all 19,573 diseases. The node2vec approach took a total of 12.0 hours to embed all nodes in the knowledge graph, including all 19,573 disease. As such, the node2vec approach is approximately 462X more efficient with respect to run time. Of note, the node2vec approach results in a Euclidean embedding for every node in the knowledge graph (which can easily be transformed to an embedding for every pair of nodes via the Haddamard product), while the DWPC approach results in an embedding only for each input pair of drug and disease. Importantly, both the DWPC feature generation and the node2vec embedding approach are both then used to train identical random forest models. Hence, the time to generate predictions is nearly identical.

**Figure 5.**
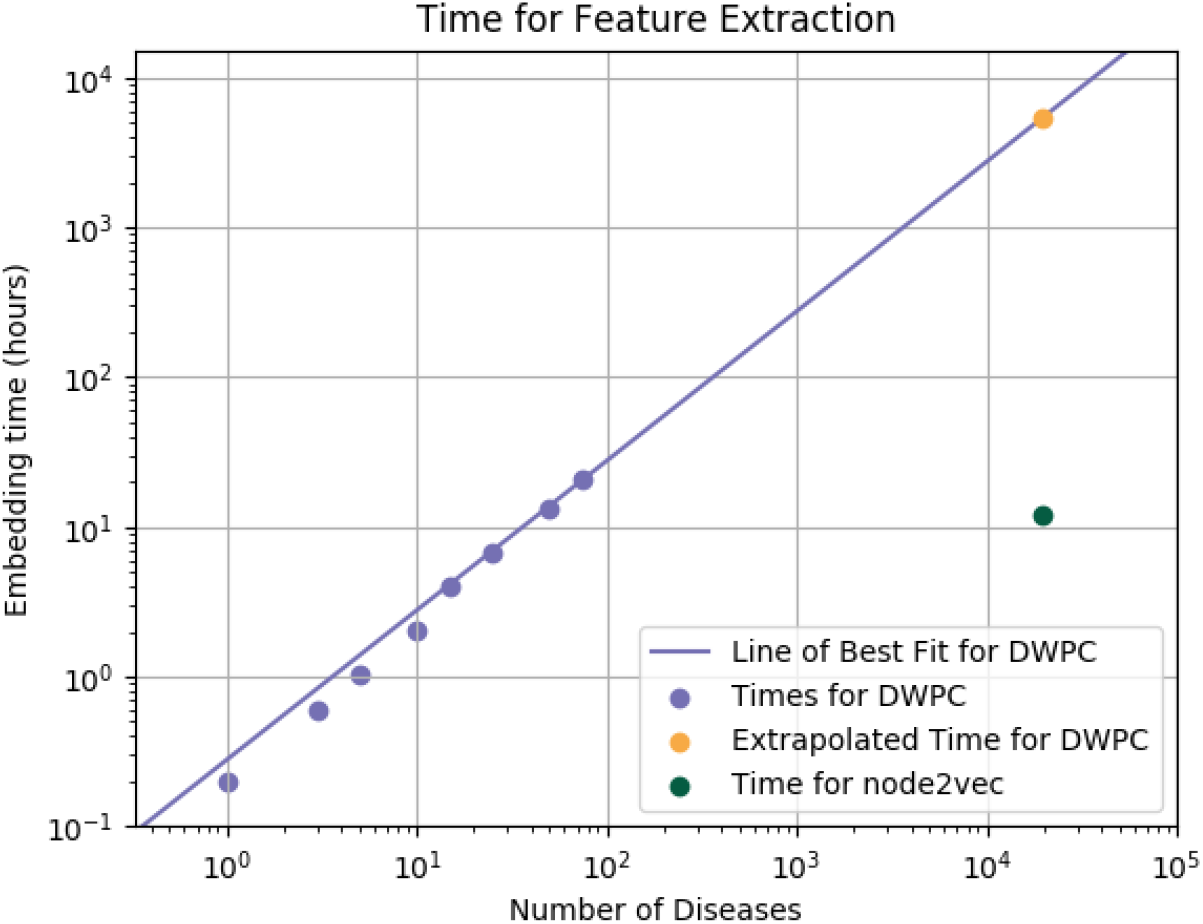
Log-log timing comparison of node embedding for both the DWPC and node2vec approaches using a fixed number (2,226) of drugs. Both methods were provided local access to the knowledge graph described in Section 2.1. The blue dots indicate the wall time of the DWPC embedding method for varying number of input diseases and the blue line indicates a linear regression fit resulting in an extrapolation to all 19,573 diseases indicated by the yellow dot. The green dot indicates the wall time of the node2vec embedding approach for all nodes in the graph (including all 19,573 diseases).

To demonstrate the potential gains to be had by the increased computational efficiency of the node2vec approach (which allows for larger training and testing set sizes), we compare in Figure 4 the performance of the DWPC approach when trained on 147 diseases to the performance of the node2vec approach using the full training data of 3,192 distinct diseases/phenotypic features. Here, the node2vec approach improves upon the DWPC approach in terms of both AUC and F1 metrics, with a mean improvement of 6.6% and 22.0% respectively.

## 5. Discussion

We have presented a novel approach to drug repurposing that leverages a graph representation of ontological data by utilizing node2vec to extract feature vectors for each node. We used this data and a random forest model to make predictions and produce probabilities for drugs treating diseases. We observed real-world examples of drug disease pairs identified as potential candidates for drug repurposing as corroborated by findings from other studies as well as clinical data from Columbia Open Health Data. We have found that the computational efficiency of this approach allows for performance improvement when compared to similar drug repurposing efforts.

Of course, careful interpretation of results and biological reality is still paramount. Despite the relatively reduced cost of drug repurposing, the cost of false positives still remains quite large given the cost of enrolling patients in clinical trials for ineffectual treatments. Thus, we think it is imperative to also take a holistic approach when examining results from this or similar methods such as when we used the plot as in Figure 3 to find a candidate threshold that filters out all but the most likely drug repurposing candidates or leveraging additional data sources such as Columbia Open Health Data [86] as in Section 3.1.

Given the continued growth in size of biological knowledge repositories, we believe it fruitful to continue to incorporate and connect publicly available repositories of biological data. Our approach demonstrates that novel discoveries may be discovered with this integrative approach. The relatively straightforward network embedding and learning method we employed provides support for this viewpoint when paired with the performance gain to be had with such an approach. In addition, with the presence of a knowledge graph, it is possible to use the predictions generated by this approach to initiate further investigation into a mechanism of action for the treatment of a disease by a given drug. Indeed, as the knowledge graph we generated contains detailed information about drug and protein target binding probabilities, biological pathways, phenotypes, etc. it may be possible for a domain expert to use this information to assess the veracity of predictions made by this approach.

We believe that there are a few key ways in which this work can be extended. First, if a well curated data set could be developed that includes not only indicated and contraindicated drug-disease pairs but also many examples of drug disease-pairs that would have no effect, performance could further be improved as the model would be less affected by publication bias of only positive results (or results of a deleterious drug interaction). Second, alternate graph embedding strategies (eg. neural network-based methods such as GraphSAGE [52] or GCN’s [61]) or alternate learning methods (eg. geometric deep learning methods [14]) may improve performance. Lastly, further addition to bioentities and relationships in the knowledge graph may provide more structure that can be leveraged by any embedding technique or learning method.

## 6. Acknowledgements

Much of the work to ingest knowledge sources into the knowledge graph described herein was carried out by the NCATS funded (see the Funding section below) RTX team at Oregon State University, including Ujjval Kumaria, Zheng Liu, Deqing Qu, Stephen Ramsey, and Yao Yao.

